# Inference about causation between body mass index and DNA methylation in blood from a twin family study

**DOI:** 10.1101/223040

**Authors:** Shuai Li, Ee Ming Wong, Minh Bui, Tuong L Nguyen, Ji-Hoon Eric Joo, Jennifer Stone, Gillian S Dite, Pierre-Antoine Dugué, Roger L Milne, Graham G Giles, Richard Saffery, Melissa C Southey, John L Hopper

## Abstract

**Background:** Several studies have reported DNA methylation in blood to be associated with body mass index (BMI), but only a few have investigated causal aspects of the association. We used a twin family design to assess this association at two life points and applied a novel analytical approach to investigate the evidence for causality.

**Methods:** The methylation profile of DNA from peripheral blood was measured for 479 Australian women (mean age 56 years) from 130 twin families. Linear regression was used to estimate the associations of methylation at ~410 000 cytosine-guanine dinucleotides (CpG), and of the average methylation at ~20 000 genes, with current BMI, BMI at age 18-21 years, and the change between the two (BMI change). A novel regression-based methodology for twins, Inference about Causation through Examination of Familial Confounding (ICE FALCON), was used to assess causation.

**Results:** At 5% false discovery rate, nine, six and 12 CpGs at 24 loci were associated with current BMI, BMI at age 18-21 years and BMI change, respectively. The average methylation of *BHLHE40* and *SOCS3* loci was associated with current BMI, and of *PHGDH* locus was associated with BMI change. From the ICE FALCON analyses with BMI as the predictor and methylation as the outcome, a woman’s methylation level was associated with her co-twin’s BMI, and the association disappeared conditioning on her own BMI, consistent with BMI causing methylation. To the contrary, using methylation as the predictor and BMI as the outcome, a woman’s BMI was not associated with her co-twin’s methylation level, consistent with methylation not causing BMI.

**Conclusion:** For middle-aged women, peripheral blood DNA methylation at several genomic locations is associated with current BMI, BMI at age 18-21 years and BMI change. Our study suggests that BMI has a causal effect on peripheral blood DNA methylation.

## Introduction

DNA methylation, that a methyl group is typically added to a cytosine-guanine dinucleotide (CpG), modifies gene expression without changing DNA sequence. Sensitive to exposures and lifestyle factors associated with health, DNA methylation has been proposed to play a critical role in the etiology of complex traits and diseases^1, 2^.

Several studies have investigated the association between DNA methylation in blood and obesity. Differences in methylation have been observed between obese and lean people^3-7^. Epigenome-wide association studies (EWAS) have reported approximately 500 CpGs to be differentially methylated in relation to body mass index (BMI)^8-16^. Methylation at some BMI-related CpGs has also been suggested to be associated with other obesity traits such as adult waist circumference^9, 10^ and BMI change^10, 16^, and obesity-related traits such as type 2 diabetes^14, 17^ and metabolic syndrome^12^.

Most of the reported associations are from cross-sectional designs, thus the causal nature of the association, i.e. whether DNA methylation has a causal effect on BMI or *vice versa*, is unknown. There is also a possibility that the observed association is due to familial confounders^18^. Mendelian randomization (MR) has been proposed as a method to assess causation^19^, and several studies have applied MR to make causal inferences between BMI and DNA methylation^8, 13, 14^.

As well as requiring knowledge and measurement of genetic variants, the validity of MR depends on certain assumptions, some of which are difficult to verify and need to be given attention^20^. Especially in the context of BMI and methylation, BMI-related genetic variants are potentially associated with DNA methylation through other obesity or metabolism pathways, which violates the assumption of no directional pleiotropy. Additionally, MR requires large-scale studies with genetic variants and DNA methylation data available for the same subjects, which might not be achievable. Obviously, other approaches for assessing causation are needed if using MR is not possible.

In this study, we aimed to investigate the association between BMI and blood DNA methylation, to replicate associations reported by previous EWAS, and to investigate the causal nature of the association using a regression-based approach to data of twins.

## Subjects and Methods

### Study sample

The sample comprised women from the Australian Mammographic Density Twins and Sisters Study (AMDTSS)^21^. A telephone-administered questionnaire was used to collect self-reported demographic information, height, current weight at interview and recalled weight at age 18-21 years. A total of 479 women, including 66 monozygotic twin (MZ) pairs, 66 dizygotic twin (DZ) pairs and 215 of their sisters from 130 families was selected ^22^. Of these, one was excluded due to her current reported BMI being an outlier. The study was approved by the Human Research Ethics Committee of the University of Melbourne. All subjects provided written informed consent.

### BMI traits

We studied three BMI traits at using measures at two ages: current BMI, BMI at age 18-21 years and BMI change between the two time-points. Current BMI and BMI at age 18-21 years were calculated using current height and the reported weight at the corresponding age.

### DNA methylation data

DNA was extracted from dried blood spots stored on Guthrie cards using a method previously described^23^. DNA was sodium bisulfite converted using the EZ DNA Methylation-Gold protocol as per manufacturers’ instructions (Zymo Research, Irvine, CA) and eluted in 20 μl elution buffer. DNA Methylation was measured using the Illumina Infinium HumanMethylation450K BeadChip (HM450) array. All laboratory work was performed at the Genetic Epidemiology Laboratory, University of Melbourne.

Raw intensity data were processed by the Bioconductor *minfi* package^24^, which included normalization of data using Illumina’s reference factor-based normalization methods (*preprocessIllumina*) and subset-quantile within array normalization (SWAN)^25^ for type I and II probe bias correction. An empirical Bayes batch-effects removal method *ComBat*^26^ was applied to minimise the technical variation across batches. All samples passed quality control. Probes with missing value (detection *P*-value >0.01) for one or more samples, with documented SNPs at the target CpG, binding to multiple locations^27^ or binding to the X chromosome, and 65 control probes were excluded, leaving 411 394 probes remaining for analysis. See Li *et al.*^22^ for more details.

### Association analyses for CpG-specific methylation

We used a linear regression model to investigate the association between CpG-specific methylation values and each BMI trait, in which the methylation M-value, the logit transformation of the percentage of methylation, was the outcome and BMI trait was the predictor. The model was adjusted for age and cell-type proportions (monocytes, B cells, natural killers, CD4+ T cells, CD+8 T cells, granulocytes) estimated using the Houseman method^28^. Parameters were estimated using the Generalised Estimating Equations (GEE) method, with family as cluster, fitted using the *geeglm()* function from the R package *geepack*. To account for multiple testing, associations with false discovery rate (FDR)^29^<0.05 were considered statistically significant.

### Association analyses for gene-average methylation

We investigated associations between gene-average methylation value and each BMI trait. We annotated CpGs to genes using the column ‘UCSC_RefGene_Name’ from the Illumina’s annotation file, resulting in a total of 19 823 genes. For each gene, the average methylation was calculated as the average Beta-value, the percentage of methylation, across CpGs annotated to that gene. The gene-average methylation was then logit transformed and the association was investigated using the same model as that for CpG analysis.

### Replication of previously reported CpGs and genes

462 CpGs and 332 genes at which there are CpGs previously reported to be associated with BMI^8-16^ were included in our study after quality control. For these CpGs and genes, we checked their associations with current BMI. For genes, the gene-average methylation was used. CpGs or genes with nominal *P*<0.05 were considered replicated.

### Causal inference analyses

We performed causal inference using Inference about Causation through Examination of FAmiliaL CONfounding (ICE FALCON), a regression-based methodology for analysing twin data^30-34^. By causal, it meant that if it were possible to vary a predictor measure experimentally then the expected value of the outcome measure would change.

As shown in Figure 1, suppose there are two variables, X and Y, measured for pairs of twins. Assume that X and Y are positively associated within an individual. Let S denote the unmeasured genetic and non-genetic factors that affect both twins; S_X_ represents those factors that influence X values only, S_Y_ those that influence Y values only, and S_XY_ those that influence both X and Y values. For the purpose of explanation, let ‘self’ refer to an individual and ‘co-twin’ refer to the individual’s twin, but recognise that these labels can be swapped and both twins within a pair are used in the analysis.

**Figure 1.**
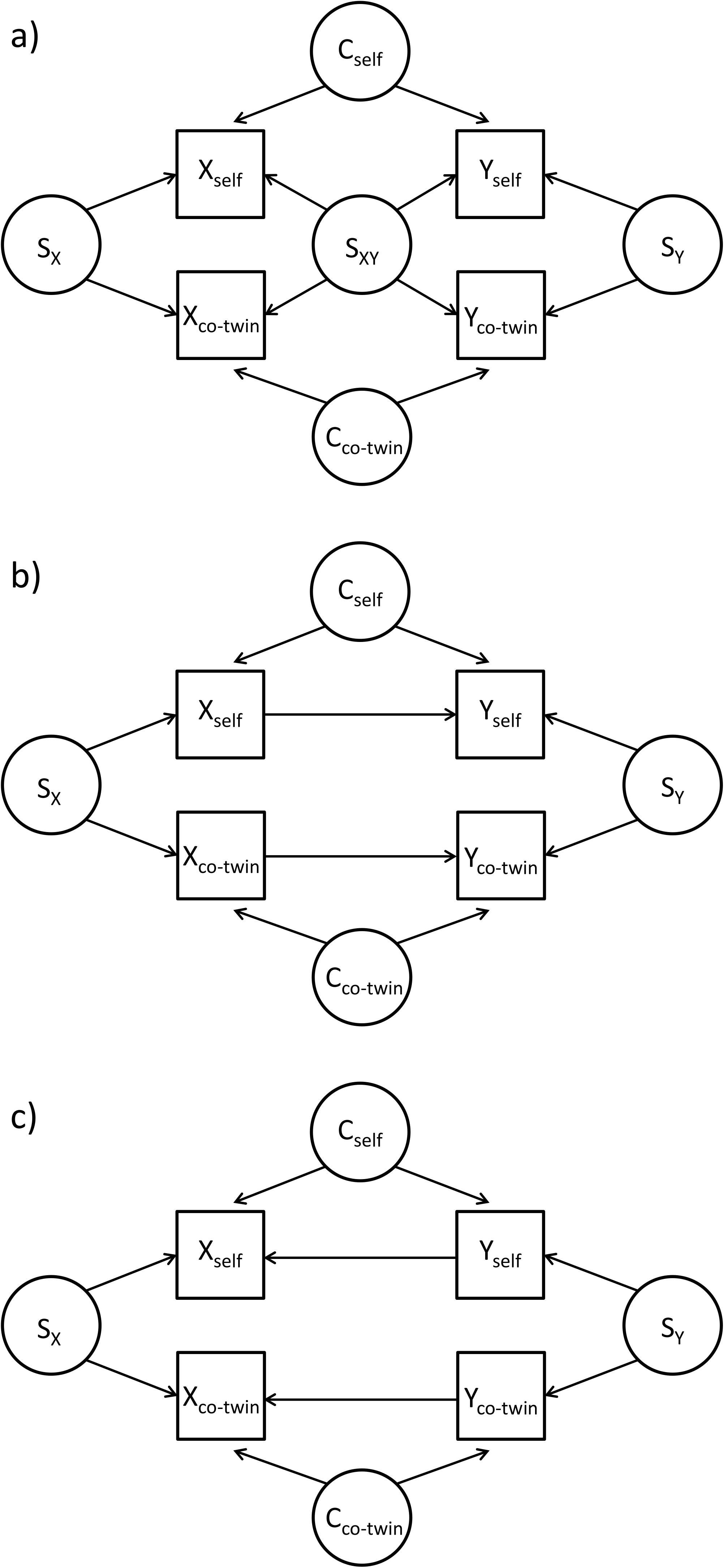
Some possible directed acyclic graphs for the causation underlying the cross-trait cross-twin correlation

If there is a correlation between Y_self_ and X_co-twin_, it might be due to a familial confounder, S_XY_ (Figure 1a). It could also be due to X having a causal effect on Y within an individual, provided X_self_ and X_co-twin_ are correlated (Figure 1b), or to Y having a casual effect on X, provided Y_self_ and Y_co-twin_ are correlated (Figure 1c). Note that the confounders specific to an individual, C_self_ and C_co-twin_, do not of themselves result in a correlation between Y_self_ and X_co-twin_.

Using GEE to take into account any correlation in Y between twins within the same pair, we fit three models:

Model 1: E(Y_self_) = α + β_self_X_self_
Model 2: E(Y_self_) = α + β_co-twin_X_co-twin_
Model 3: E(Y_self_) = α + β’_self_X_self_ + β’_co-twin_X_co-twin_

If the correlation between Y_self_ and X_co-twin_ is solely due to familial confounders (Figure 1a), the marginal association between Y_self_ and X_self_ (β_self_ in Model 1) and the marginal association between Y_self_ and X_co-twin_ (β_co-twin_ in Model 2) must both be non-zero. Adjusting for X_self_, however, the conditional association between Y_self_ and X_co-twin_ (β’_co-twin_ in Model 3) is expected to attenuate from β_co-twin_ in Model 2 towards the null. Similarly, adjusting for X_co-twin_ (Model 3), the conditional association between Y_self_ and X_co-twin_ (β’_self_ in Model 3) is expected to attenuate from β_self_ towards the null.

If the correlation between Y_self_ and X_co-twin_ is solely due to a causal effect from X to Y (Figure 1b), by adjusting for X_self_ the pathway between these two variables is blocked and the conditional association (β’_co-twin_ in Model 3) is expected to be null. Adjusting for X_co-twin_ will have no influence on the pathway between Y_self_ and X_self_, so the conditional association β’_self_ in Model 3 is expected to be β_self_ in Model 2.

If the correlation between Y_self_ and X_co-twin_ is solely due to a causal effect from Y to X (Figure 1c), the pathway through X_self_ is blocked due to X_self_ as a collider, and the pathway through SY is blocked due to that GEE analysis in effect conditions on SY, so there is no marginal association between Y_self_ and X_co-twin_, and β_co-twin_ of Model 2 is expected to be zero.

We first applied ICE FALCON to methylation at CpGs reported by Wahl *et al.*, the majority of which has been suggested to be consequential to BMI by a bidirectional MR analysis^14^. We analysed a methylation score based on the 77 CpGs with *P*<0.05 in our study. The methylation score was calculated as the sum of the products of the methylation Beta-value and the reported effect size of each CpG. The data for MZ pairs were used. The models were adjusted for age and cell-type proportions as what was done in association analyses. We first used the methylation score to be Y and current BMI to be X and regressed the methylation score on current BMI. We then swapped X and Y to regress current BMI on the methylation score and undertook the same analyses. We made statistical inference about the changes in regression coefficients by bootstrapping. That is, twin pairs were randomly sampled with replacement to generate 1,000 new datasets with the same sample size as the original dataset. ICE FALCON was then applied to each dataset to calculate the changes in regression coefficients for that dataset.

We then applied ICE FALCON to methylation at CpGs and genes identified in our study. For each BMI trait, we similarly investigated the methylation at CpGs as a methylation score. For a locus containing multiple CpGs, only the CpG with the smallest *P*-value was included in the methylation score. BMI at age 18-21 years and it related methylation were not investigated, given that the methylation score was not correlated within MZ twin pairs (r = -0.07, 95% CI: -0.31, 0.17).

## Results

### Characteristics of the sample

For the women included in the analytic sample, the mean (standard deviation [SD]) age was 56.4 (7.9) years. The mean (SD) current BMI, BMI at age 18-21 years and BMI change was 26.8 (5.7), 21.1 (3.3) and 5.6 (5.0) kg/m^2^, respectively. Current BMI was positively correlated with BMI at age 18-21 years (r = 0.49, 95% CI: 0.42, 0.55), and more strongly with BMI change (r = 0.81, 95% CI: 0.78, 0.84), and BMI at age 18-21 years was weakly negatively associated with BMI change (r = -0.11, 95% CI: -0.20, -0.02).

### Associations with CpG-specific methylation

Methylation at nine, six and 12 CpGs was found to be associated with current BMI, BMI at age 18-21 years and BMI change, respectively (Table 1). The genomic inflation factor λ was 1.10, 1.03 and 1.04 for current BMI, BMI at age 18-21 years and BMI change, respectively (Q-Q plots in Supplementary Figure 1; Manhattan plots in Supplementary Figures 2-4). Methylation at two CpGs (cg12992827 and cg00636368) was negatively associated with both current BMI and BMI change. CpGs cg12992827^10, 14^, cg18181703^12-14, 16^, cg09349128^10, 13, 14^, and the *NOD2* locus^13, 14, 16^ have been reported previously. The other CpGs or loci have not been reported before.

**Table 1.**
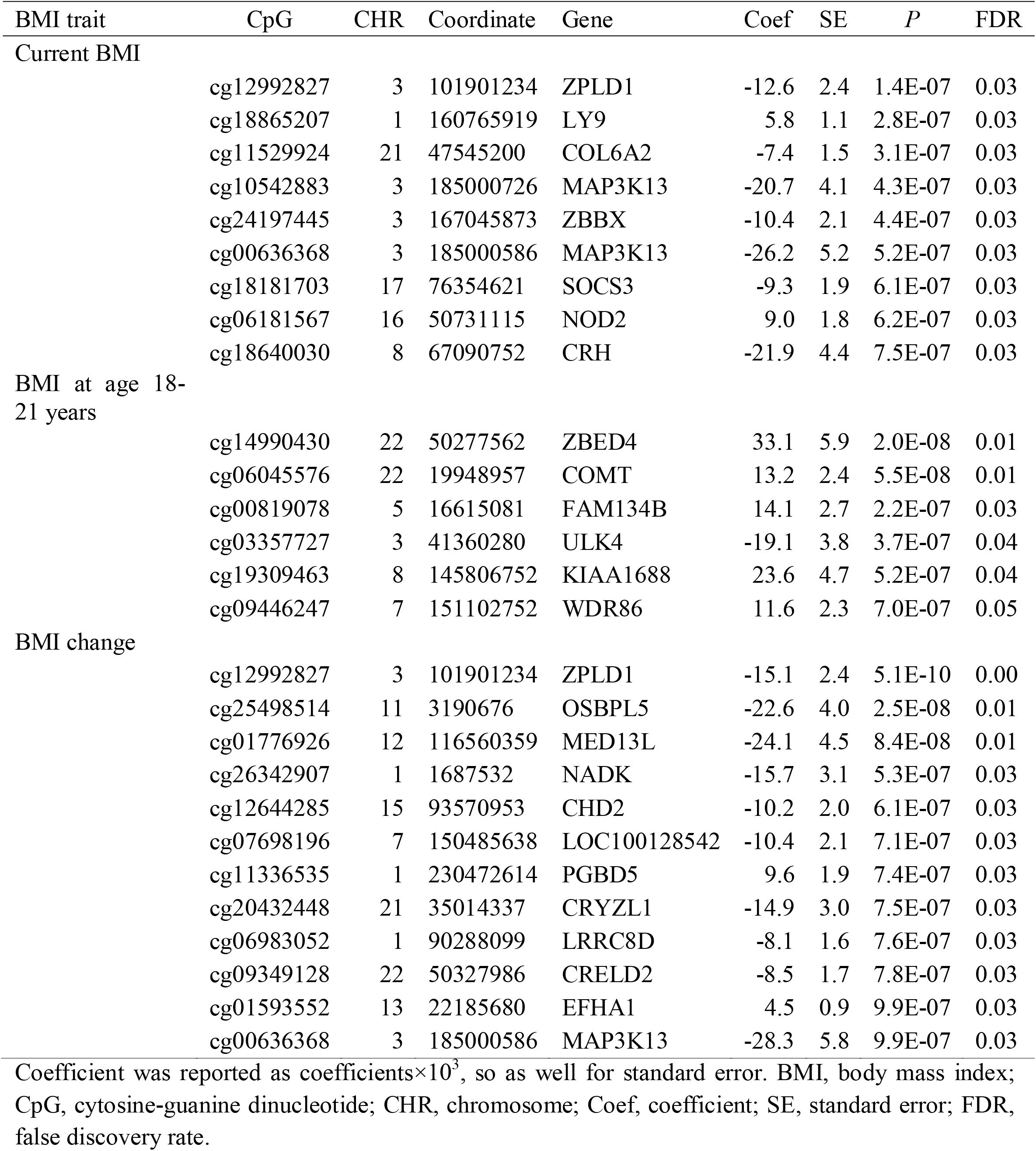
CpGs at which methylation was found to be associated with the three BMI traits

Supplementary Table 1 shows the associations between methylation at the identified CpGs for one BMI trait and the other BMI traits. At nominal *P*<0.05, methylation at all CpGs for current BMI was associated BMI change in the same direction, and *vice versa*. Methylation at approximately 50% CpGs for current BMI was associated with BMI at age 18-21 years in the same direction, and *vice versa*. None of CpGs for BMI change at which methylation was associated with BMI at age 18-21 years, nor *vice versa*. Supplementary Table 2 shows the associations between methylation at the identified CpGs and corresponding BMI trait adjusting for the other BMI traits. At nominal *P*<0.05, except for a few CpGs, methylation at all the identified CpGs was associated with corresponding BMI trait independent of the other BMI traits.

### Associations with gene-average methylation

The average methylation at *BHLHE40* and *SOCS3* were both negatively associated with current BMI (*BHLHE40*: coefficient[coef]=-1.1×10^−3^, 95% CI: -1.5, -0.7, *P*=3.8×10^−7^, FDR=0.01; *SOCS3*: coef= -1.6×10^−3^, 95% CI: -2.3, -1.0, *P*=1.4×10^−6^, FDR=0.01), and the average methylation at *PHGDH* was negatively associated with BMI change (coef=-2.2×10^−3^, 95% CI: -3.1, -1.3, *P*=2×10^−6^, FDR=0.04). No gene was found to be associated with BMI at age 18-21 years. The genomic inflation factor λ was 1.13, 0.95 and 1.04 for current BMI, BMI at age 18-21 years and BMI change, respectively. (Q-Q plots in Supplementary Figure 5). *SOCS3*^12-15^ and *PHGDHt*^9, 12-15^ have been previously reported, and *BHLHE40* has not been reported before. Methylation at seven, five and five CpGs annotated to *BHLHE40, SOCS3* and *PHGDH* were associated with their respective BMI trait with nominal *P*<0.05, respectively (Supplementary Figures 6-8).

At nominal *P*<0.05, the average methylation at *BHLHE40* and *SOCS3* was also associated with BMI at age 18-21 years and BMI change. The average methylation at *PHGDH* was also associated with current BMI but not with BMI at age 18-21 years. The average methylation at *BHLHE40* and *SOCS3* was associated with current BMI independent of BMI at age 18-21 years and BMI change, and the average methylation at *PHGDH* was associated with BMI change independent of current BMI (data not shown).

### Replication of previously reported associations

Altogether, 151 CpG associations were replicated with nominal *P*<0.05 and were in the same direction as that from previous studies, and the 59 most significant CpGs also had FDR<0.05. Our study reported the first replication for 104 CpGs (Supplementary Table 3). 41 gene associations were replicated with nominal *P*<0.05, and the six most significant genes also had FDR<0.05 (Supplementary Table 4).

### Causal inference analyses results

The ICE FALCON results for methylation at CpGs reported by Wahl *et al.^14^* are shown in Table 2. From the analyses with BMI was the predictor and the methylation score the outcome, a woman’s methylation score was associated with her own BMI (Model 1; β_self_ = 0.96, 95% CI: 0.45, 1.47) and with her co-twin’s BMI (Model 2; β_co-twin_= 0.57, 95% CI: 0.06, 1.08). Conditioning on her co-twin’s BMI (Model 3), β’_self_ remained unchanged (*P*=0.93), while conditioning on her own BMI (Model 3), β’_co-twin_ attenuated by 95.5% (95CI: 7.0%, 181.2%; *P*=0.03) to be 0.03 (95% CI: -0.50, 0.55). From the analyses with the methylation score was the predictor and BMI the outcome, a woman’s BMI was associated with her own methylation score (Model 1; β_self_ = 0.78, 95% CI: 0.14, 1.41) but not with her co-twin’s methylation score (Model 2; β_co-twin_= -0.02, 95% CI: -0.33, 0.30). In Model 3, β’_self_ remained unchanged (*P*=0.16) compared with β_self_ in Model 1, and β’_co-twin_ increased by 2500.0% (95%CI: 622.2%, 5044.3%; *P*=0.03) compared with β_co-twin_ in Model 2 to be 0.38 (95% CI: -0.01, 0.77). These results are consistent with the findings by Wahl *et al*. that methylation at these CpGs are consequential to BMI.

**Table 2.** Results from the ICE FALCON analyses for CpG-specific methylation*

| CpG | Predictor | Coefficient | Model 1 |  | Model 2 |  | Model 3 |  | Change |  |
| --- | --- | --- | --- | --- | --- | --- | --- | --- | --- | --- |
|  |  |  | Est (SE) | *P* | Est (SE) | *P* | Est (SE) | *P* | Est (SE) | *P* |
| CpGs reported by Wahl *et al.* |  |  |  |  |  |  |  |  |  |  |
| | BMI | $\beta_{\text{self}}$ | 0.96 (0.26) | 2.3E-04 | | | 0.95 (0.31) | 2.3E-03 | -0.01 (0.16) | 9.3E-01 |
| | | $\beta_{\text{co-twin}}$ | | | 0.57 (0.26) | 2.8E-02 | 0.03 (0.27) | 9.2E-01 | -0.54 (0.26) | 3.3E-02 |
| | Methylation score | $\beta_{\text{self}}$ | 0.78 (0.32) | 1.6E-02 | | | 1.00 (0.32) | 1.6E-03 | 0.22 (0.16) | 1.6E-01 |
| | | $\beta_{\text{co-twin}}$ | | | -0.02 (0.16) | 9.2E-01 | 0.38 (0.20) | 5.4E-02 | 0.40 (0.18) | 3.0E-02 |
| Current BMI CpGs |  |  |  |  |  |  |  |  |  |  |
| | BMI | $\beta_{\text{self}}$ | 0.40 (0.06) | 7.5E-13 | | | 0.45 (0.07) | 2.3E-09 | 0.05 (0.05) | 3.6E-01 |
| | | $\beta_{\text{co-twin}}$ | | | 0.28 (0.07) | 9.4E-05 | -0.07 (0.07) | 3.3E-01 | -0.35 (0.08) | 3.6E-05 |
| | Methylation score | $\beta_{\text{self}}$ | 4.57 (1.10) | 3.3E-05 | | | 5.91 (1.12) | 1.2E-07 | 1.33 (0.67) | 4.6E-02 |
| | | $\beta_{\text{co-twin}}$ | | | -0.79 (0.78) | 3.1E-01 | 2.20 (0.78) | 5.0E-03 | 3.00 (0.98) | 2.2E-03 |
| BMI change CpGs |  |  |  |  |  |  |  |  |  |  |
| | BMI change | $\beta_{\text{self}}$ | 0.57 (0.07) | 2.0E-16 | | | 0.49 (0.09) | 1.2E-07 | -0.08 (0.08) | 3.2E-01 |
| | | $\beta_{\text{co-twin}}$ | | | 0.47 (0.09) | 6.3E-08 | 0.12 (0.10) | 2.0E-01 | -0.35 (0.09) | 3.9E-05 |
| | Methylation score | $\beta_{\text{self}}$ | 3.83 (0.78) | 8.5E-07 | | | 4.25 (0.70) | 1.5E-09 | 0.43 (0.44) | 3.3E-01 |
| | | $\beta_{\text{co-twin}}$ | | | 0.26 (0.61) | 6.7E-01 | 1.41 (0.62) | 2.3E-02 | 1.15 (0.56) | 3.9E-02 |

ICE FALCON, inference about causation through examination of familial confounding; CpG, cytosine-guanine dinucleotide; BMI, body mass index; Est, estimate; SE, standard error.
\*Regression coefficient from the analyses in which methylation score as the predictor was reported as coefficient × 10, so as well for standard error.

Similarly, the results for methylation at CpGs identified in our study are consistent with that current BMI and BMI change have a causal effect on methylation at their related CpGs, respectively (Table 2). The same causation was inferred for gene-average methylation (Supplementary Table 5).

## Discussion

We performed EWASs between BMI at two life points and blood DNA methylation, and found that methylation at several loci was associated with BMI at middle age, BMI at age 18-21 years and BMI change. Methylation at some of these loci, *ZPLD1*^10, 14^, *SOCS3*^12-16^, *CRELD2*^10, 13, 14^, *NOD2*^13, 14, 16^ and *PHGDH*^9, 13, 14, 16^, has been previously reported to be associated with BMI. Associations at the other loci, such as *LY9*, *COL6A2*, *ZBBX*, *MAP3K13*, *CRH*, etc., do not appear to have been previously reported. We replicated the associations for methylation at 151 CpGs and 41 genes reported by previous studies, and reported the first replication for 104 of the CpGs. The investigation of causation suggests that BMI and BMI change have a causal effect on methylation, but not *vice versa*.

Several of the identified loci are involved in inflammatory pathways, which are biologically relevant to obesity. *LY9* (lymphocyte antigen 9) is involved in regulation of the development of thymic innate memory-like CD8+ T cell^35^ and interleukin-17 production^36^. *MAP3K13* (mitogen-activated protein kinase kinase kinase 13) is involved in activation of mitogen-activated protein kinases (MAPK)^37^. It has also been found to regulate NF-κB transcription factor activity^38^. *NOD2* (nucleotide binding oligomerization domain containing 2) is involved in immune response by playing roles in several biological pathways, such as MAPK activation^39^, cytokine production^40-42^ and NF-κB activation^43, 44^. *SOCS3* (suppressor of cytokine signalling 3) plays a role in regulation of inflammatory response by negatively regulating cytokine signalling. It is also involved in the Gene Ontology (GO) pathway for negative regulation of insulin receptor signalling (GO:0046627). *CRH* (corticotropin releasing hormone) is a member of the corticotropin-releasing factor family. It is included in the GO pathway for inflammatory response (GO:0006954). Previous studies have also reported methylation at loci involved in inflammatory pathways to be associated with BMI^9, 10, 12-16^.

Some of the identified loci are related to other pathways. Three loci are involved in oxidation reduction process. *NADK* (NAD kinase) is a NAD+ kinase, which catalyses the reduction process from NAD to NADP. *CRYZL1* (crystallin zeta like 1) encodes quinone oxidoreductase and is involved in oxidation reduction process. *PHGDH* (phosphoglycerate dehydrogenase) is a member of oxidoreductase family. Two loci are involved in serine metabolism. *ULK4* (unc-51 like kinase 4) is a serine/threonine kinase. *PHGDH* is involved in the early steps of L-serine synthesis. Associations between methylation at loci related to serine metabolism and BMI have also been reported by previous studies^9, 13, 16^.

Our study implies that BMI at different life stages influences DNA methylation through different pathways. For example, we found evidence that the loci at which methylation was associated with BMI at age 18-21 years differed from those with current BMI, and especially with BMI change. Though current BMI was associated with methylation at the loci of BMI change and *vice versa*, the associations with current BMI and BMI change are independent of each other, suggesting roles for different pathways.

Our study provides evidence for causation underlying the BMI-methylation association. We replicated that BMI has a causal effect on methylation at the CpGs reported by Wahl *et* al.^14^. We found evidence consistent with that the differential methylation at our identified CpGs is secondary to BMI, but not *vice versa*. Findings from previous studies^8, 13, 45^ also support such a causal relationship. Inflammation is proposed to play a role in the development of some obesity-related diseases, and several of the previously reported and our discovered loci are involved in inflammatory pathways, so our findings imply that methylation mediates the effects of obesity on obesity-related diseases. More research on this mediating role of methylation is warranted.

Our study demonstrates how ICE FALCON can be useful for the investigation of causation underlying epigenetic associations. ICE FALCON shares similarities with a bidirectional MR approach. In the scenario represented by the Figure 1b, we essentially use SX as an instrumental variable for X_self_. However, S_X_ is not measured, so we use X_co-twin_ as a surrogate instrumental variable (Similarly, Y_co-twin_ is used as a surrogate for S_Y_, the instrumental variable for Y_self_, when the roles of Y and X are swapped). S_X_ includes all causes of familial correlation in X that are specific to X, which is stronger than a finite number of genetic variants that are assumed to be specific to X, thus the results from ICE FALCON are less biased by the strength of the instrumental variable. S_X_, by definition, is not associated with confounders of the relationship between X and Y within an individual and has no directional pleiotropy (any association or directional pleiotropy is captured in S_XY_). Even though the surrogate instrumental variable X_co-twin_ is associated with the confounders or has directional pleiotropy, ICE FALCON still works. For example, in a scenario where the association between X_co-twin_ and Y_self_ is mediated not only through S_X_ and X_self_ but also through other unmeasured factors such as familial confounders, i.e., a mixture of Figures 1a and 1b, even though X_co-twin_ will still be associated with Y_self_ after adjusting for X_self_, β_co-twin_ is still expected to attenuate towards the null, given that adjusting for X_self_ blocks the pathway X_co-twin_ ← S_X_ → X_self_ → Y_self_. ICE FALCON can be an alternative method for investigating causation when genetic variants data are unavailable, or the assumption underlying MR analyses are suspect. ICE FALCON can also be used as an independent method to replicate the findings from MR analyses, and *vice versa*. However, requirements such as the need for substantial within-pair correlations may limit the application of ICE FALCON, which was the case for BMI at age 18-21 years in our study.

Our gene-average methylation approach is of potential utility for analysing EWAS. Such utility is supported to some extent by our replication of previously reported genes including *SOCS3* and *PHGDH*. A CpG-by-CpG approach is typically used in EWAS. However, this approach requires a stringent genome-wide significance level, e.g., 10^−7^ when half a million of CpGs are analysed and the Bonferroni adjustment is applied, which may not be achievable. The gene-based approach aimed to address this challenge by reducing the number of tests to 20 000. This approach can potentially find novel associations missed by the CpG-by-CpG approach, such as the association for *BHLHE40*.

Our study has several strengths. We studied three BMI traits at different life points. We used a novel family-based analytical approach to investigate the causation underlying the cross-sectional association and reached the same conclusion as the MR approach. We investigated gene-average methylation, which can potentially provide evidence for novel associations.

One limitation of our study is that BMI traits were self-reported, thus are subject to bias. Another limitation is that we studied methylation in peripheral blood, not in tissues more relevant to adiposity such as adipose tissue. However, cross-tissue investigations suggest moderate to good overall consistency between blood and adiposity-relevant tissues in the context of BMI-methylation association. In the study by Demerath *et al.*, associations for 64% of CpGs identified in blood were replicated in adipose tissue^10^. In the study by Wahl *et al.*, blood methylation levels at BMI-related CpGs had moderate to high correlations with methylation levels in adiposity-relevant tissues^14^. More importantly, given that peripheral blood samples are easily accessible, any association with BMI detectable in this sample is potentially valuable for large-scale public health translation on population level.

In conclusion, we found evidence that in peripheral blood from middle aged women, DNA methylation at several loci is associated with current BMI, BMI at age 18-21 years and BMI change between the two ages. Additionally, our study found evidence consistent with BMI having a causal effect on peripheral blood DNA methylation.

## Acknowledgements

We would like to thank all women participating in this study. The data analysis was facilitated by Spartan, the High Performance Computer and Cloud hybrid system of the University of Melbourne.

The AMDTSS was facilitated through the Australian Twin Registry, a national research resource in part supported by a Centre for Research Excellence Grant from the National Health and Medical Research Council (NHMRC) APP 1079102. The AMDTSS was supported by NHMRC (grant numbers 1050561 and 1079102), Cancer Australia and National Breast Cancer Foundation (grant number 509307).

SL is supported by the Australian Government Research Training Program Scholarship from the University of Melbourne. TLN is supported by a NHMRC Post-Graduate Scholarship and the Richard Lovell Travelling Scholarship from the University of Melbourne. MCS is a NHMRC Senior Research Fellow. JLH is a NHMRC Senior Principal Research Fellow.

## Conflict of Interest

The authors declare no conflicts of interest.

